# SPATIALLY PATTERNED PODOCYTE STATE TRANSITIONS COORDINATE AGING OF THE GLOMERULUS

**DOI:** 10.64898/2026.02.27.708213

**Authors:** Christopher Chaney, Jeffrey W. Pippin, Uyen Tran, Diana G. Eng, Juliang Wang, Thomas Carroll, Stuart J. Shankland, Oliver Wessely

## Abstract

**Background:** With the US population living longer, the risk, incidence, prevalence and severity for chronic kidney diseases become more abundant. Glomerular diseases are the leading cause for chronic and end-stage kidney disease. Yet, the cellular responses and the underlying mechanisms of progressive glomerular disease, which ultimately leads to glomerulosclerosis and loss of kidney function with advancing age, are poorly understood.

**Methods:** Kidneys of young (4 months-old), middle-aged (20 months-old) and aged (24 months-old) mice were separated into outer cortex and juxta-medullary region and processed for single nuclei transcriptomics. Focusing on the aging glomerulus data were analyzed using a state-of-the-art analysis pipeline dissecting out the cellular age- and kidney region-specific responses.

**Results:** Global analysis of the transcriptome reveals regional-specific differences that are detectable across multiple cell types exemplified by the expression of Napsa as a *bona-fide* juxta-medullary marker. In contrast aging led to rather cell type-specific responses. In the glomerulus, healthy podocytes were characterized by expression of canonical podocyte genes; conversely the senescent, aged podocytes were characterized by the down-regulation of canonical podocyte genes and the emergence of inflammatory and senescent signatures. Interestingly, these senescent podocytes were primarily located in the juxtamedullary region suggesting that juxtamedullary podocytes are more sensitive. Yet, instead of aging being defined by distinct cell states, the profiles, as well as ligand-receptor and pseudotime analyses suggest that podocytes aging is selective and coordinated, not universal degeneration. This was different to the other glomerular cell types, parietal epithelial cells, glomerular endothelial cells and mesangial cells. While they also as existed in different subpopulations, they exhibited little regional-, or age-depended changes. Finally proximal tubular aging manifested itself as discrete cellular states.

**Conclusions:** The single nuclei transcriptomics of the aging kidney provides a mechanistic explanation for regional susceptibility of nephrons and suggests that the future therapeutic strategies need to consider the cellular and spatial complexity of the glomerulus.

## INTRODUCTION

The global population is aging. According to the US census bureau the numbers of Americans ages 65 and older will more than double over the next 40 years.^1^ Similarly, in Europe older individuals will account for nearly a third of the population by 2050.^2^ In the kidney, numerous studies have shown that glomerular and tubulointerstitial diseases are disproportionately more common in people over 65 years old,^3,4^ with worse outcomes compared to younger individuals with the same disease.^5,6^ Chronic kidney disease (CKD) and end-stage kidney disease (ESKD) are also disproportionately increased in the elderly,^7^ with patients undergoing chronic dialysis for the first time now being predominantly older adults.^8–10^ In a recent “healthy adult study”, of those older than 70 years, 48% had glomerulosclerosis and 47% had CKD.^11^ Yet, the significance of these observations is still controversial and it is still undefined what constitutes normal kidney aging and what is disease.^12,13^ Aging of the glomerulus in general and the podocytes in particular has been described.^13,14^ Glomerular Filtration Rate (GFR) declines after age 40 by 0.8-1.0% per year,^15,16^ and kidneys from healthy donors aged 70-75 years have 48% fewer intact nephrons compared to those of younger donors aged 19-29 years.^17^ This observation is consistent with an estimated annual loss of 6,000-6,500 nephrons after age 30.^17,18^

The healthy aging kidney is characterized by progressive glomerular and tubulointerstitial fibrosis,^14,19–21^ accompanied by cellular changes in the tubulointerstitium, the vasculature and the glomerulus.^14,22,23^ For example, our group has reported age-associated changes to podocytes,^24–30^ glomerular parietal epithelial cells (PECs),^24,30,31^ and glomerular endothelial cells (GEnCs).^24,30^ Landmark studies have identified the depletion of glomerular podocytes as a sentinel pathophysiological lesion in the development of age-related glomerulosclerosis.^32–40^ Aged podocytes undergo cellular stress,^41^ senescence^24^ and exhibit reduced function including decreased secretion of VEGF.^42^ Podocyte depletion is particularly harmful because these terminally differentiated epithelial cells are an integral component of the glomerular filtration barrier do not proliferate and regenerate only in a limited fashion. ^43–52^

Although podocyte depletion is considered a central lesion in age-related glomerulosclerosis, this paradigm does not explain why structurally intact glomeruli exhibit functional decline, why aging affects multiple glomerular cell types simultaneously, or why susceptibility differs between cortical and juxtamedullary nephrons. These observations suggest that aging may reflect coordinated changes within the glomerular cellular network rather than failure of a single cell population.

However, most prior studies have characterized structural and cellular phenotypes, whereas the molecular programs operating within individual glomerular cell types during normal aging remain incompletely defined. In particular, aging may involve coordinated changes across multiple glomerular cell populations mediated through intercellular signaling networks [ADD REF: cell–cell communication in kidney aging or microenvironmental regulation].

Importantly, not all nephrons age equivalently. Glomeruli located in the outer cortex and those in the juxtamedullary region differ in filtration pressure, blood flow, oxygenation, and metabolic demand.^53–56^ These regional differences have been associated with differential susceptibility to injury and sclerosis,^57–60^ yet the molecular basis for this heterogeneity during normal aging is unknown.

Because aging is cell-type specific, spatially heterogeneous, and potentially dependent on intercellular signaling, approaches that resolve transcriptional states at single-cell resolution are required. To address these fundamental biological questions, we performed single-nucleus RNA sequencing (snRNA-seq) of mouse kidneys separated into outer cortex and juxtamedullary regions, comparing healthy young, middle-aged, and elderly mice. We hypothesized that aging of the glomerulus is driven by coordinated, cell-type–specific transcriptional programs and altered intercellular signaling that differ between outer cortical and juxtamedullary regions.

## MATERIAL AND METHODS

### Nuclei Isolation, Library Preparation, and Sequencing

Single-nucleus RNA sequencing (snRNA-seq) was performed on mouse kidneys collected at 4, 20, and 25 months of age. Kidneys were micro-dissected into outer cortical (OC) and juxtamedullary (JM) regions prior to nuclei isolation. For each age–region combination, six technical replicates were processed independently. Libraries were prepared using the 10x Genomics Chromium platform and sequenced according to the manufacturer’s protocol.

### Primary Processing, Quality Control, and Normalization

Sequencing data were processed using established single-cell RNA-seq workflows in R and Bioconductor. ^61,62^ Empty droplets were removed and per-nucleus quality control metrics were calculated, including mitochondrial transcript proportion and library complexity using DropletUtils^63^ and scater^64^. Low-quality nuclei were excluded based on outlier detection criteria. Counts were normalized using deconvolution-based size factors and log-transformed expression values were generated using scran^65^ and scuttle.^66^ Genes exhibiting biological variability were identified and used for downstream dimensionality reduction.

### Dataset Integration and Latent Representation Learning

To enable joint analysis across samples and sequencing batches, datasets were converted to AnnData format using zellkonverter.^67^ Data were analyzed in Python using Scanpy.^68^ Batch correction and dimensionality reduction were performed using the scVI deep generative modeling framework^69^ to learn a shared latent representation of transcriptional state across all nuclei. The learned latent space served as the basis for all downstream embedding, clustering, and comparative analyses.

### Global Cell State Identification and Annotation

A neighborhood graph was constructed from the scVI latent space and used to generate a low-dimensional embedding using UMAP^70^ and community structure using Louvain clustering.^71^ Clusters were annotated using canonical marker genes and expert curation to define major renal cell populations. This global atlas provided the reference framework for lineage-specific analyses.

### Marker Gene Identification

Marker genes were identified using the EIGEN framework^72^ on stratified subsets of nuclei to ensure balanced representation across age and anatomical zone. Analyses were performed both at the cluster level and after assignment of cell type identities. Rare and unclassified populations were excluded from lineage-level marker discovery.

### Lineage-Specific Subpopulation Analyses

To characterize aging-associated heterogeneity within individual glomerular and tubular lineages, each major cell type was reanalyzed independently using a shared analytical strategy. For each lineage, nuclei were embedded in a local manifold, clustered to identify subpopulations, and subjected to marker discovery. Parameters were optimized per lineage and are provided in Supplementary Methods. For glomerular endothelial and mesangial populations, a two-stage approach was used to first isolate glomerular subsets and then resolve finer substructure. For large populations such as proximal tubule cells, representative subsampling was performed prior to analysis.

### Functional Gene Program Analysis

Gene set activity was quantified using AUCell^73^ with curated mouse gene sets from MSigDB.^74^ Gene programs associated with cluster identity, age, and anatomical region were identified by applying EIGEN^72^ to nucleus-level gene set activity scores. For selected analyses, diffusion-based imputation was used to enhance detection of coordinated gene programs based on MAGIC methodology.^75^

### Trajectory Inference

Developmental relationships among podocyte subpopulations were inferred using pseudotime trajectory analysis implemented in slingshot^76^. No trajectory inference was performed for other cell types.

### Reproducibility and Software Availability

All analyses were conducted using reproducible R and Python pipelines. Detailed software versions, parameters, and configuration settings are provided in the Supplementary Methods. Processed data and analysis code will be made publicly available upon publication.

### Machine Learning Model to Identify Kidney Regions

Region prediction was performed on an initial set of 9,255 representing from the OC, JM and the medulla (M). We used the XGBoost R package to train a gradient boosting machines (GBM) model to assign a nuclei/cell to the OC, JM and the M. Gradient boosting successively trained an ensemble of shallow decision trees to do classification. Each decision tree tried to correct the mistakes made by the previous trees. In classification, all trees then perform a weighted majority vote to determine spatial labels. We randomly split all our cells into training set (80% of all cells, 7,404) and testing set (1851). We used the 7,404 cells and their corresponding spatial labels (OC, JM, M) and trained a GBM model (parameters: learning rate 0.001, max tree depth 5, gamma = 3). Prediction performance was evaluated on the 1,851 cells not used in the training process. The xgb.plot.importance function was used to identify genes that contribute to the OC vs. JM vs. M separation.

## RESULTS

### Age- and Region-Specific Single Nuclei Transcriptomics of Mouse Kidneys

To investigate how aging alters glomerular cell states, we performed single-nucleus transcriptomic profiling of mouse kidneys at 4, 20, and 25 months of age, corresponding approximately to human ages of 20, 60, and 80 years. To capture both spatial and temporal variation during aging, kidneys were microdissected into outer cortex (OC) and juxtamedullary (JM) compartments (**Figure 1A**) prior to processing for single-nucleus mRNA sequencing (snRNA-seq) following KPMP protocols.^77^ After quality control and integration, the dataset comprised 890,093 high-quality nuclei (mean 13,375 reads per nucleus; median 3,378 UMIs and 1,959 detected genes). Unsupervised clustering identified 11 major renal cell populations, annotated using established marker genes (**Figure 1B,C**; **Table 1**). Regional composition of the data matched expected nephron anatomy: collecting duct and distal tubular populations were enriched in OC samples, whereas loops of Henle were enriched in JM samples (**Figure 1E**). In contrast, cell type proportions were largely preserved across ages (**Figure 1D**), indicating that aging does not primarily alter kidney cellular composition but instead influences transcriptional state within existing populations, establishing a framework to examine lineage-specific and spatially patterned aging responses.

**Figure 1.**
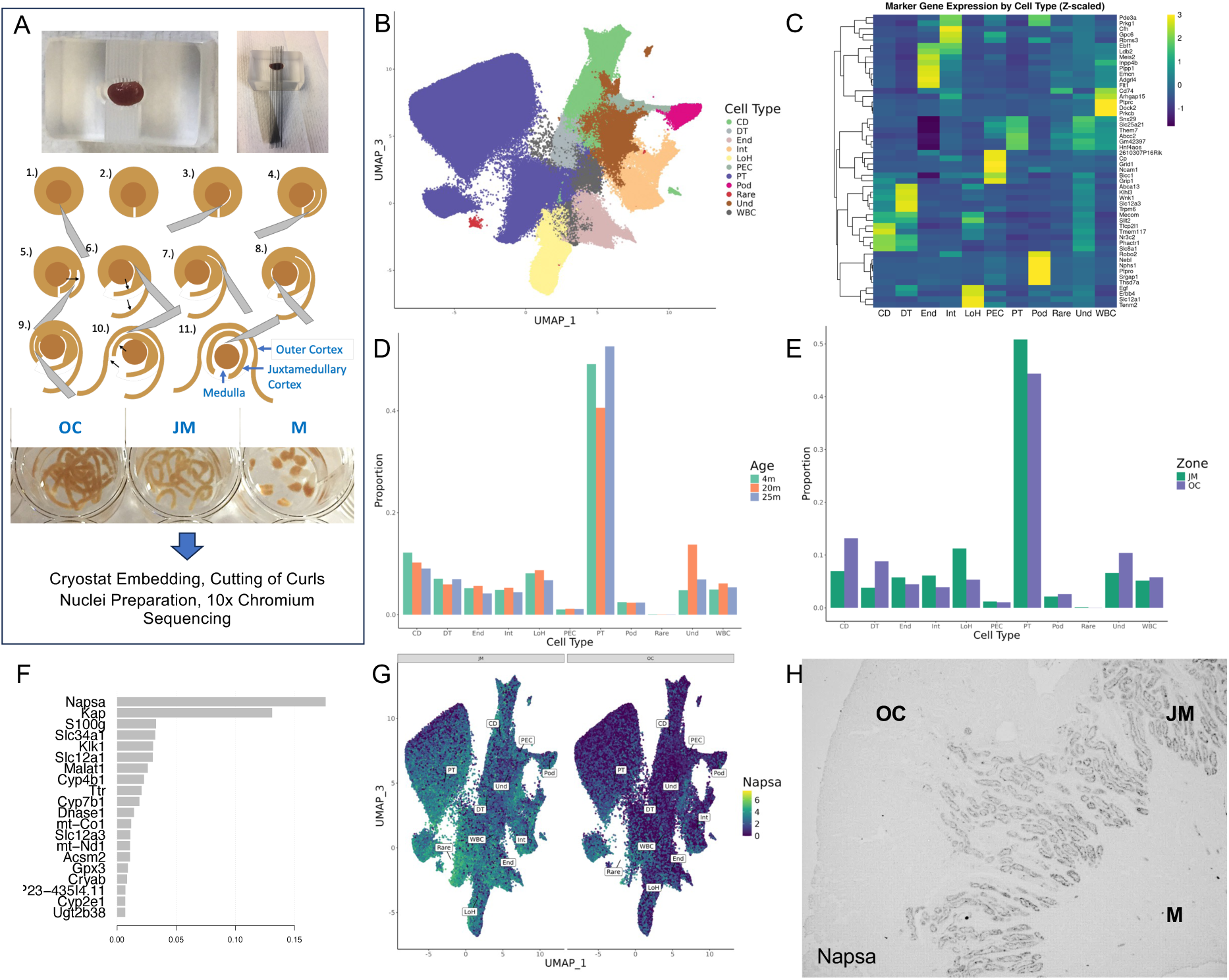
Overview of Experimental Design and Cell Type Identification. (**A**) To dissect the cellular composition of the kidney in young (4 months), middle-aged (20 months) and aged (24 months) mice, isolated kidneys were cryo-sectioned and separated by region (OC - outer cortex, JM - juxtamedullary region and M - medulla) following the 11 steps as illustrated. Regional dissections were then processed for snRNA-seq following the protocol established by the KPMP consortium.^77^ (**B,C**) Following quality control and doublet removal, 890,093 nuclei were retained for analysis. UMAP visualization (B) shows nuclei colored by annotated cell type based on canonical marker expression identified with the EIGEN framework. This yielded collecting duct (CD), distal tubule (DT), endothelium (End), interstitium (Int), Loops of Henle (LoH), parietal epithelium (PEC), proximal tubule (PT), podocyte (Pod) as well as a rare (<0.1% of nuclei, Rare), and an undetermined (Und) population. Each major cell type is characterized by top marker genes as depicted in the heatmap (**C**) showing scaled expression levels (z-scores). (**D,E**) Bar graphs show a relatively even distribution of nuclei across cell types parsed by age (D) and region (E). Data are expressed as the proportion of nuclei from each age group. (**F-H**) Comparing the transcriptomic data based on location identified those genes that contribute most to the separation between OC vs. JM cells (F). UMAP projections of nuclei separated by JM and OC zones and overlayed by the log-normalized expression of *Napsa* mRNA demonstrate higher expression in JM-derived nuclei (G). Consistent with the transcriptomic data immunohistochemistry confirms elevated Napsa protein levels in the JM, but not the OC or the M (H).

**Table 1:**
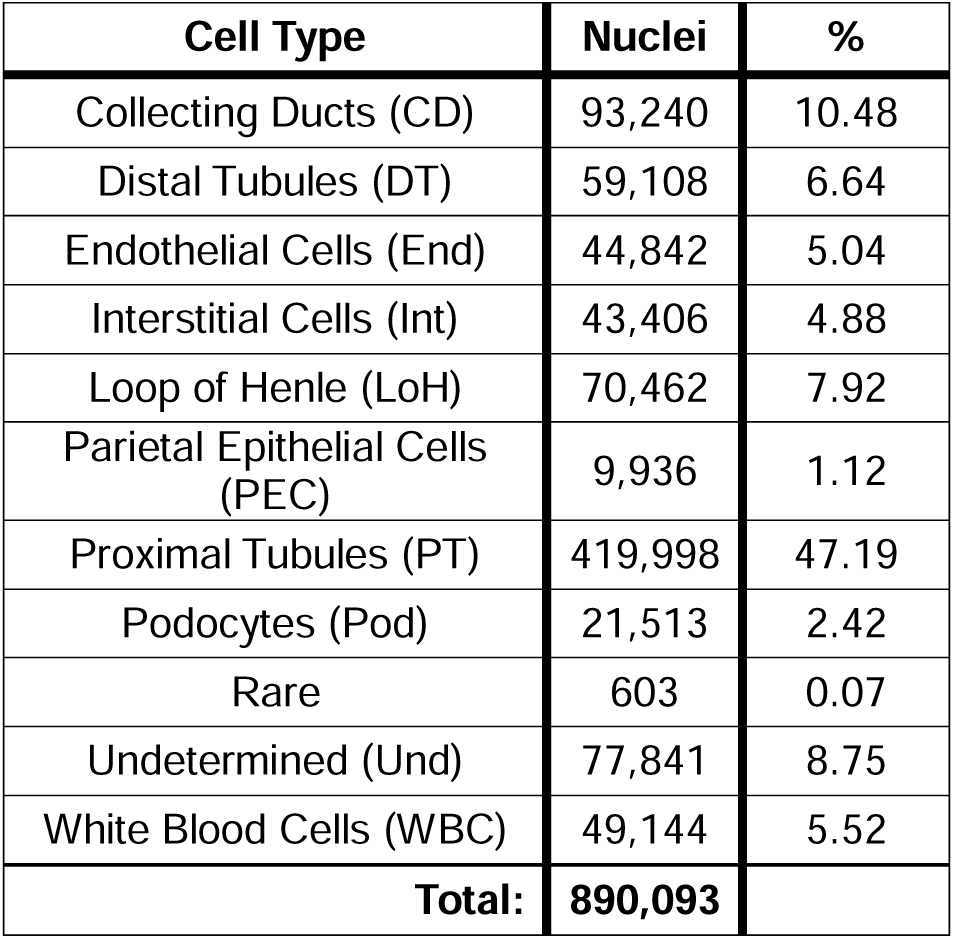
Distribution of the nuclei to the different cell types.

| Cell Type | Nuclei | % |
| --- | --- | --- |
| Collecting Ducts (CD) | 93,240 | 10.48 |
| Distal Tubules (DT) | 59,108 | 6.64 |
| Endothelial Cells (End) | 44,842 | 5.04 |
| Interstitial Cells (Int) | 43,406 | 4.88 |
| Loop of Henle (LoH) | 70,462 | 7.92 |
| Parietal Epithelial Cells (PEC) | 9,936 | 1.12 |
| Proximal Tubules (PT) | 419,998 | 47.19 |
| Podocytes (Pod) | 21,513 | 2.42 |
| Rare | 603 | 0.07 |
| Undetermined (Und) | 77,841 | 8.75 |
| White Blood Cells (WBC) | 49,144 | 5.52 |
| **Total:** | **890,093** |  |

We next asked whether transcriptional information alone could distinguish nuclei originating from OC and JM regions independent of cell type. To address this, we developed a classifier that predicts the anatomical origin of individual nuclei based solely on gene expression (**Supplemental Figure S1A–F**). The model identified a small set of highly discriminative genes, among which Napsin A Aspartic Peptidase (Napsa) and Kidney androgen regulated protein (Kap) were the strongest predictors (**Figure 1F**). Kap is known to be enriched in proximal tubules of the kidney,^78,79^ whereas comparatively little is known about Napsa beyond its association with cancer and expression in lung and kidney.^80^ Feature plots demonstrated cell type–independent enrichment of Napsa expression in JM-derived nuclei (**Figure 1G**). This spatial pattern was confirmed by immunohistochemistry in mouse kidneys (**Figure 1H**) and in human kidney tissue (**Supplemental Figure S1G**).

Together, these findings are consistent with under-reported kidney region-specific transcriptional identities that are detectable across multiple cell types. This establishes a spatial context as an intrinsic component of renal cellular state rather than solely an anatomical classification.

### snRNA-seq Analysis of Proximal Tubules

To determine how aging affects specific renal lineages, we first examined the proximal tubule compartment, which has been extensively characterized in development and injury.^81–84^ A total of 419,998 nuclei were annotated as proximal tubule cells. After subsampling and re-clustering, eight transcriptionally distinct proximal tubule subpopulations were identified (**Figure 2A,B**). Segment identity was resolved using canonical markers. The proximal tubule transcription factor HNF4g was enriched in clusters 2, 3, and 6 (**Figure 2C**). Clusters 4 and 5 expressed the S1 marker Slc5a12 (**Figure 2D**), whereas cluster 6 expressed the S2/S3 marker Slc9a8 (**Figure 2E**). These molecular assignments aligned with anatomical distribution: clusters 2–5 were enriched in the OC, while cluster 6 was enriched in the JM (**Figure 2G,H**). In contrast, clusters 1, 7, and 8 lacked many canonical proximal tubule markers and instead expressed injury-associated genes including Fau (**Figure 2B,F**). These populations were strongly enriched in the JM (**Figure 2G,H**). Although individual clusters showed variable representation across ages, no single cluster expanded monotonically with aging (**Figure 2I,J**). However, gene set enrichment analysis revealed that all three clusters were enriched for aging and senescence signatures (Rodwell_Kidney_Aging_Up, Rodwell_Kidney_Aging_Down, Fridman_Senescence_Up, and Reactome_Senescence_Associated_Secretory_Pathway). These findings demonstrate that aging manifests as discrete cellular states rather than uniform degeneration across a lineage, consistent with the PT_VCAM1 population described in mouse and human kidneys.^82,83^

**Figure 2.**
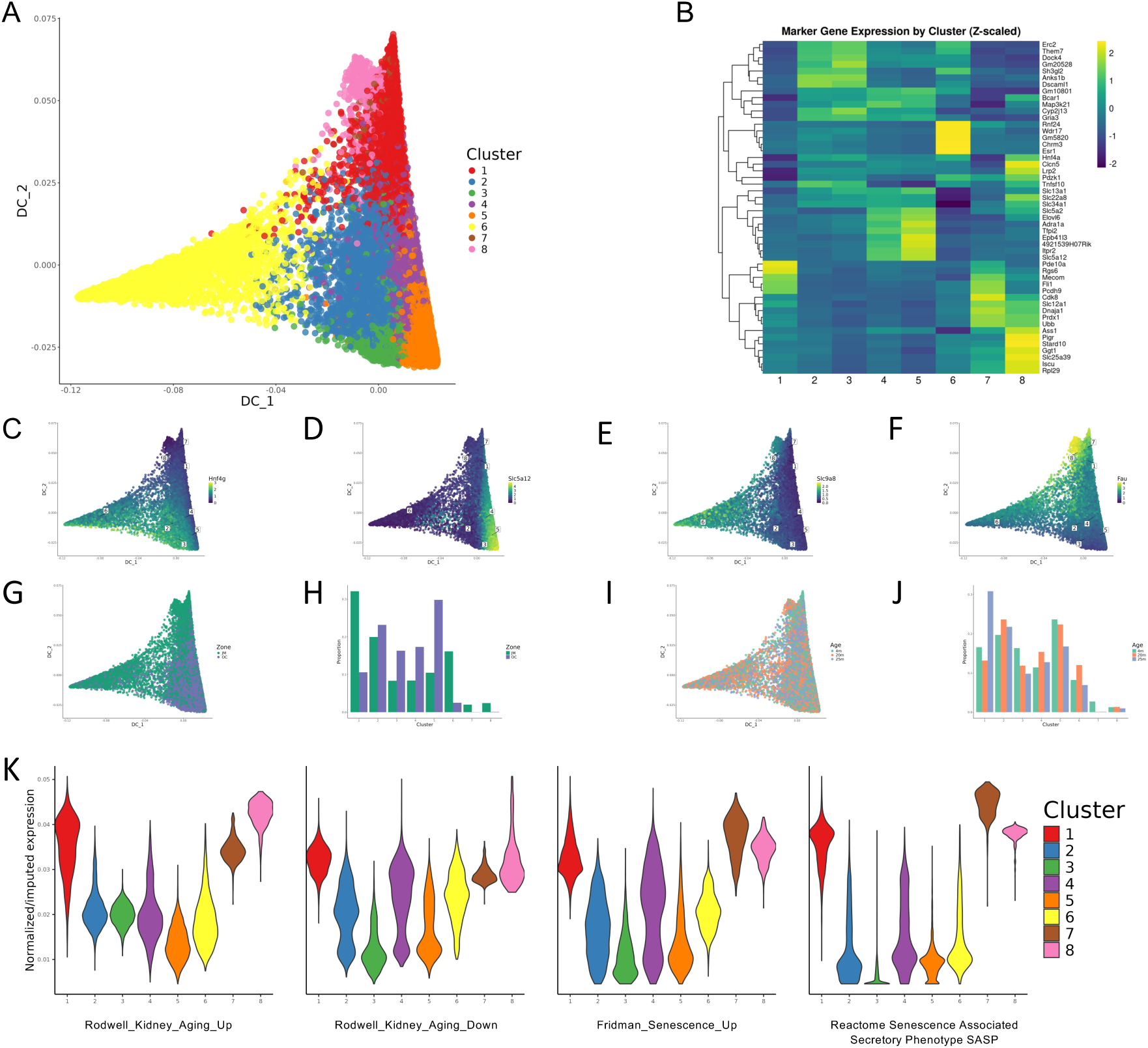
Transcriptional Heterogeneity of Proximal Tubule Cells Across Age and Anatomical Zone. (**A**) Sub-clustering of proximal tubule cells and their visualization by diffusion map yields a total of 8 clusters with six of them representing terminal regions (clusters 1, 2, 3, 5, 6, and 8), and cluster 4 and 7 likely being transitional states. Note that cluster 2 is projecting slightly out of the plane shown and can be better seen in 3-D view (see in Supplemental Movie S1). (**B**) Heatmap showing the top marker genes characteristic for each of the individual proximal tubular clusters with the expression values being normalized within the proximal tubule subset (B). (**C–F**) Gene-overlayed diffusion maps show characteristic marker genes expressed in selected clusters, the proximal tubular transcription factor *HNF4G* (C), the proximal tubule S1 segment marker *SLC5A12* (D), the proximal tubule S2/3 segment marker *SLC9A8* (E) and the apoptosis regulator *Fau* (F) highlighting transcriptional diversity across the proximal tubules. (**G,H**) Diffusion map colored by anatomical zone (G) and the corresponding bar graph (H) show regional specific distributions with an enrichment of JM-derived nuclei in clusters 1, 6, 7, and 8 and OC-derived nuclei in clusters 3,4, and 5. (**I,J**) Diffusion map colored by age (I) and the corresponding bar graph (J) show enrichment relatively even distribution of age across the different proximal tubular clusters with the exception of cluster 1, which is enriched for nuclei from of the 25-month-old mice and cluster 7, which nearly exclusively contains nuclei of the young 4-months old mice. (**K**) Violin plots graphing the presence of cells enriched for aging/senescence signature gene sets (Rodwell_Kidney_Aging_Up, Rodwell_Kidney_Aging_Down, Fridman_Senescence_Up, and Reactome_Senescence_Associated_Secretory_Pathway) showing enrichment in clusters 1, 7, and 8 suggesting that these clusters are impacted by age or injury.

### Podocyte Aging Reflects Spatially Biased Transcriptional State Transitions

We next examined glomerular podocytes known to exhibit an age-dependent decline ^14,40,85,86^ and regional differences between OC and JM podocytes.^87–89^ The dataset contained 21,513 podocyte nuclei. Bulk comparison of canonical podocyte gene programs revealed reduced expression of structural and differentiation-associated genes with age, particularly in the JM (**Supplemental Figure S3A**), whereas senescence and inflammatory programs were increased in 25-month kidneys (**Supplemental Figure S3B**). To resolve cellular heterogeneity, podocytes were subclustered, revealing a tetrahedral diffusion manifold composed of nine transcriptionally distinct states (**Figure 3A**; **Supplemental Movie S1**). The vertices corresponded to discrete podocyte populations (**Figure 3B**). Cluster 0 (red circles in **Figure 3A**) represented a canonical differentiated state marked by *Wt1*, *Mafb*, *Nphs1*, *Podxl*, and *Synpo* (**Figure 3B,C**); other vertices represented alternative transcriptional states including *Bnc2*-expressing and *Cyp7b1*-enriched populations (**Figure 3B,D,E**). The last of the vertex populations, cluster 5 (orange circles in **Figure 3B**) expressed genes such as *Camk1d* and *Tmsb4x* and was strongly enriched for aging, senescence, and inflammatory pathways (RODWELL_AGING_KIDNEY_NO_BLOOD_UP, FRIDMAN_SENESCENCE_UP, REACTOME_SENESCENCE_ASSOCIATED_SECRETORY_PHENOTYPE_SASP, and GOBP_POSITIVE_REGULATION_OF_ACUTE_INFLAMMATORY_RESPONSE), identifying a senescent podocyte state (**Figure 3G–J**). Notably, this population was almost exclusively composed of juxtamedullary podocytes and did not progressively expand with chronological age (**Figure 3K,L**), indicating that senescence-associated remodeling is spatially restricted rather than purely time-dependent.

**Figure 3.**
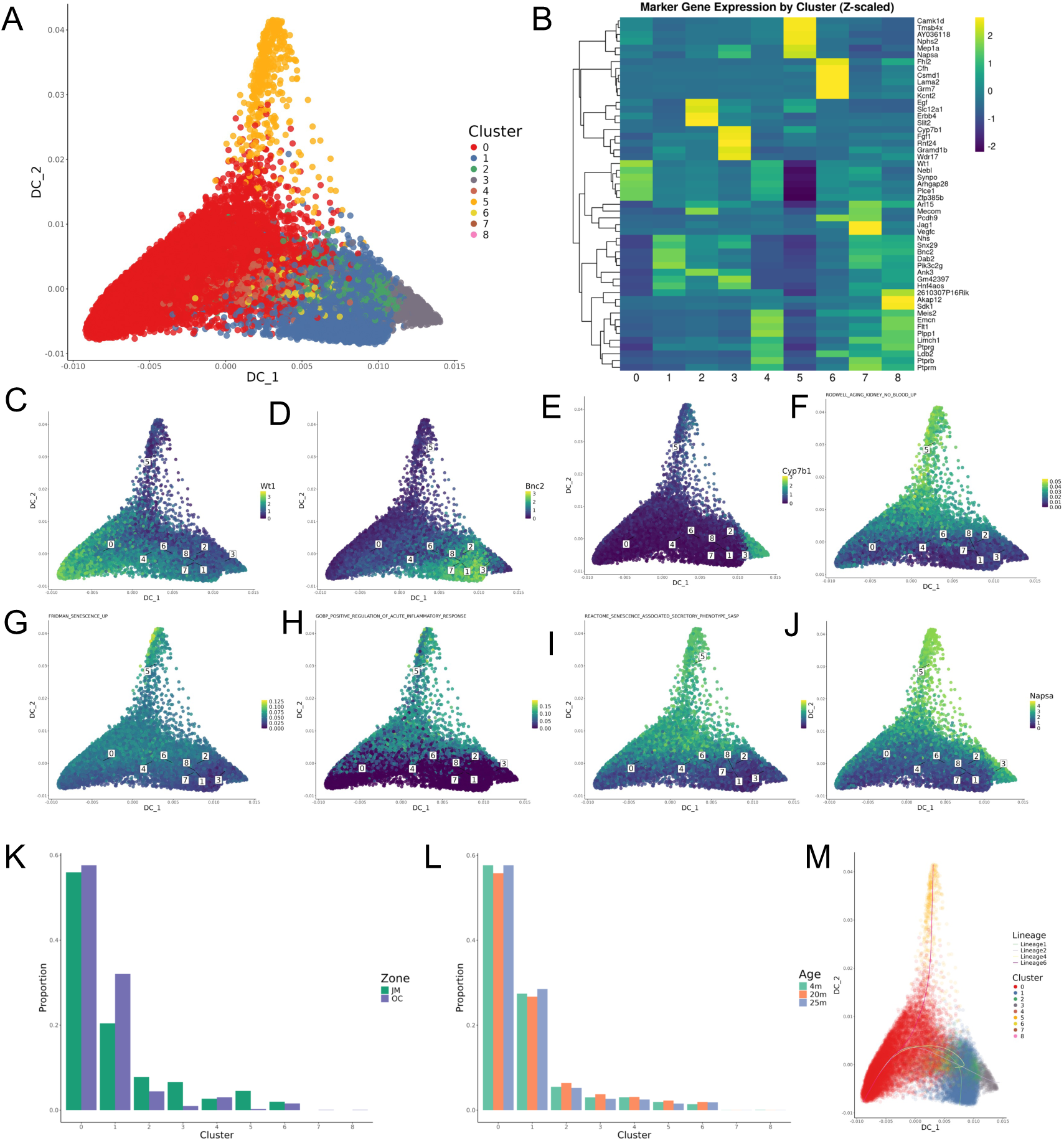
Podocyte Heterogeneity is Shaped More by Anatomical Zone than by Age. (**A-E**) Diffusion map of sub-clustered podocytes (A) yields 9 populations that are arranged in a tetrahedron-like pattern with 4 vertices (see also 3-D view in Supplemental Movie S2). The heatmap (B) shows for each podocyte cluster the imputed expression of top marker genes, which were identified by the EIGEN algorithm and visualized by z-scores. Overlay diffusion maps showing expression of representative marker genes for three of the four vertices, the canonical podocyte marker *Wt1* (C), Basonuclin-2 (*Bnc2*) (D) and the oxysterol 7-alpha-hydroxylase (*Cyp7b1*) (E). The 4^th^ vertex is positive for cell exhibiting high enrichment scores for the following aging/senescence signature gene sets, i.e., Rodwell_Kidney_Aging_Up (F), Fridman_Senescence_Up (G), Reactome_Senescence_Associated_Secretory_Pathway (H), and GOBP_Positive_Regulation_of_Acute_Inflammatory_Response (I). (**J,K**). Regional differences in podocyte sub-clusters can be seen when the diffusion map is overlayed with the JM-marker *Napsa*, which is enriched in clusters 2, 3, and 5 (J) as well as bar plot showing anatomical zone distribution across podocyte clusters (K). Note that cluster 1 is enriched in the JM nuclei. (**L**) Bar plot for the age distribution across the podocyte clusters shows a relatively even distribution of nuclei from all age groups across all clusters. (**M**) Diffusion map with Slingshot-inferred trajectories colored by lineage highlights potential transitions among podocyte identifies a path from healthy podocytes to senescent ones (lineage 6, pink), as well as to/from the other vertices (lineages 1, 2 and 4).

Consistent with this organization, multiple podocyte states exhibited regional bias: cluster 1 was enriched in OC podocytes, and regional identity across states could be predicted by *Napsa* expression (**Figure 3F,K,L**), supporting the existence of a regionally encoded podocyte transcriptional program. Intermediate clusters (2, 4, 6, and 7) showed reduced canonical podocyte gene expression and increased expression of transcriptional regulators such as *Meis2* and *Mecom* (**Figure 3A,B**), suggesting transitional states. Trajectory inference using slingshot^76^ identified a direct transition from the canonical state (cluster 0) to the senescent state (cluster 5), while alternative paths traversed intermediate populations (**Figure 3M**).

Together, these findings indicate that podocytes occupy a structured continuum of transcriptional states, including a spatially restricted path to senescence and additional regionally biased intermediate trajectories. Thus, podocyte aging reflects coordinated state transitions within a cellular network rather than a uniform degenerative process.

### Non-podocyte Glomerular Cell Types Exhibit Relative Transcriptional Stability During Aging

We next examined the remaining glomerular cell populations: parietal epithelial cells (PECs), glomerular endothelial cells (GEnCs), and mesangial cells. A total of 9,936 <u>PEC</u> nuclei were extracted from the global dataset and sub-clustering identified eight transcriptionally distinct PEC states (**Figure 4A,B**). Most PEC subpopulations were similarly represented between OC and the JM, with only modest regional bias (cluster 3 enriched in OC; cluster 7 enriched in JM) (**Figure 4C**; **Supplemental Figure S4A**). Importantly, none of the PEC populations demonstrated consistent age-dependent expansion or depletion (**Figure 4D**; **Supplemental Figure S4B**).

**Figure 4.**
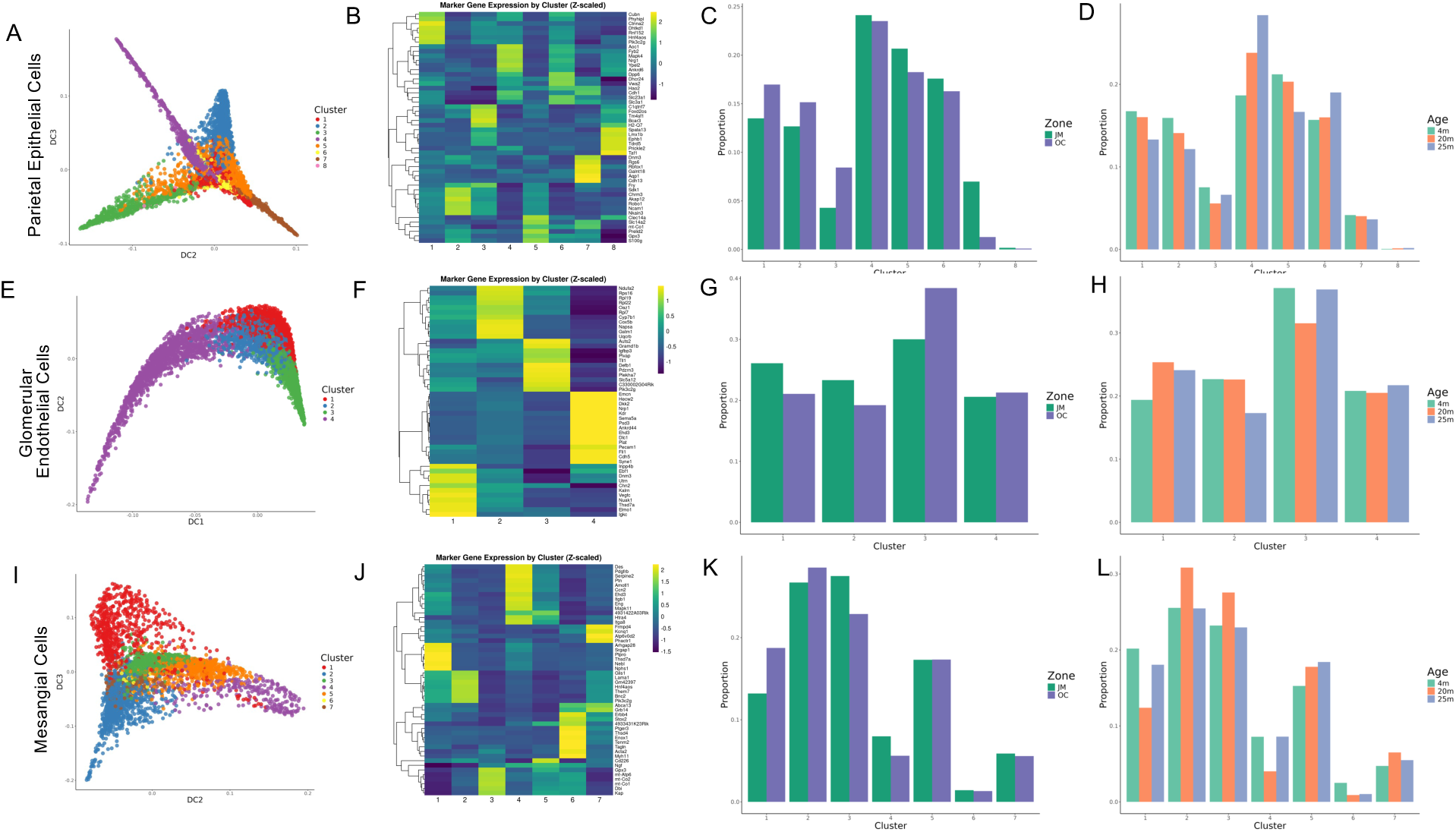
Transcriptional Heterogeneity of Non-Podocyte Glomerular Cell Types. (**A-D**) Sub-clustering of the parietal epithelial cells (PECs) identifies 8 clusters that are well-separated in the diffusion map (A) and are characterized by unique set of top marker genes in the heatmap (B). Bar plot for the regional distribution of the PECs across the clusters shows a relatively even distribution except for cluster 3 being enriched in the OC and cluster 7 enriched in the JM (C). Conversely, there was no difference in age distribution across the PEC clusters (D). (**E-H**) Sub-clustering of glomerular endothelial cells (GEnCs) identifies 4 clusters with unique marker gene expression that are relatively evenly distribute both in respect to OC *vs.* JM as well as age. (**I-L**) Sub-clustering of the mesangial cells identifies 7 clusters that are well-separated in the diffusion map (I), characterized by unique set of top marker genes (M), which show little differences in distribution, when parsed by region (K) or age (L). For all three cell types, the smooth transitions between clusters suggest continuous cell state variation.

<u>GEnCs</u> were extracted from the endothelial cluster (**Figure 1B,C**) using *Itga1*^90^ and resolved into four transcriptionally distinct subpopulations (**Figure 4E,F**). In contrast to podocytes, these endothelial states showed no detectable enrichment by region or age (**Figure 4G,H**; **Supplemental Figure S4C,D**).

<u>Mesangial cells</u> were similarly isolated from the interstitial cluster based on *Sgk1*^90^ (**Figure 1B,C**) and segregated into seven transcriptionally distinct subpopulations (**Figure 4I,J**). As with GEnCs, mesangial populations were largely preserved across both anatomical region and age (**Figure 4K,L**; **Supplemental Figure S4E,F**), with only a modest reduction of cluster 6 in older animals.

Together, these findings indicate that aging selectively remodels specific glomerular lineages - podocytes - within a shared microenvironment rather than uniformly affecting all glomerular cells.

### Podocytes Act as Signaling Hubs within the Glomerulus

To determine whether podocyte state changes could influence the surrounding glomerular microenvironment, we performed ligand–receptor analysis comparing ligands expressed by podocytes with corresponding receptors across glomerular cell types (podocytes, PECs, GEnCs, and mesangial cells). Interactions were evaluated across anatomical location (OC vs. JM) and age (4, 20, and 25 months). The resulting interaction map revealed structured and cell-type-specific communication networks rather than uniform signaling (**Figure 5**).

**Figure 5.**
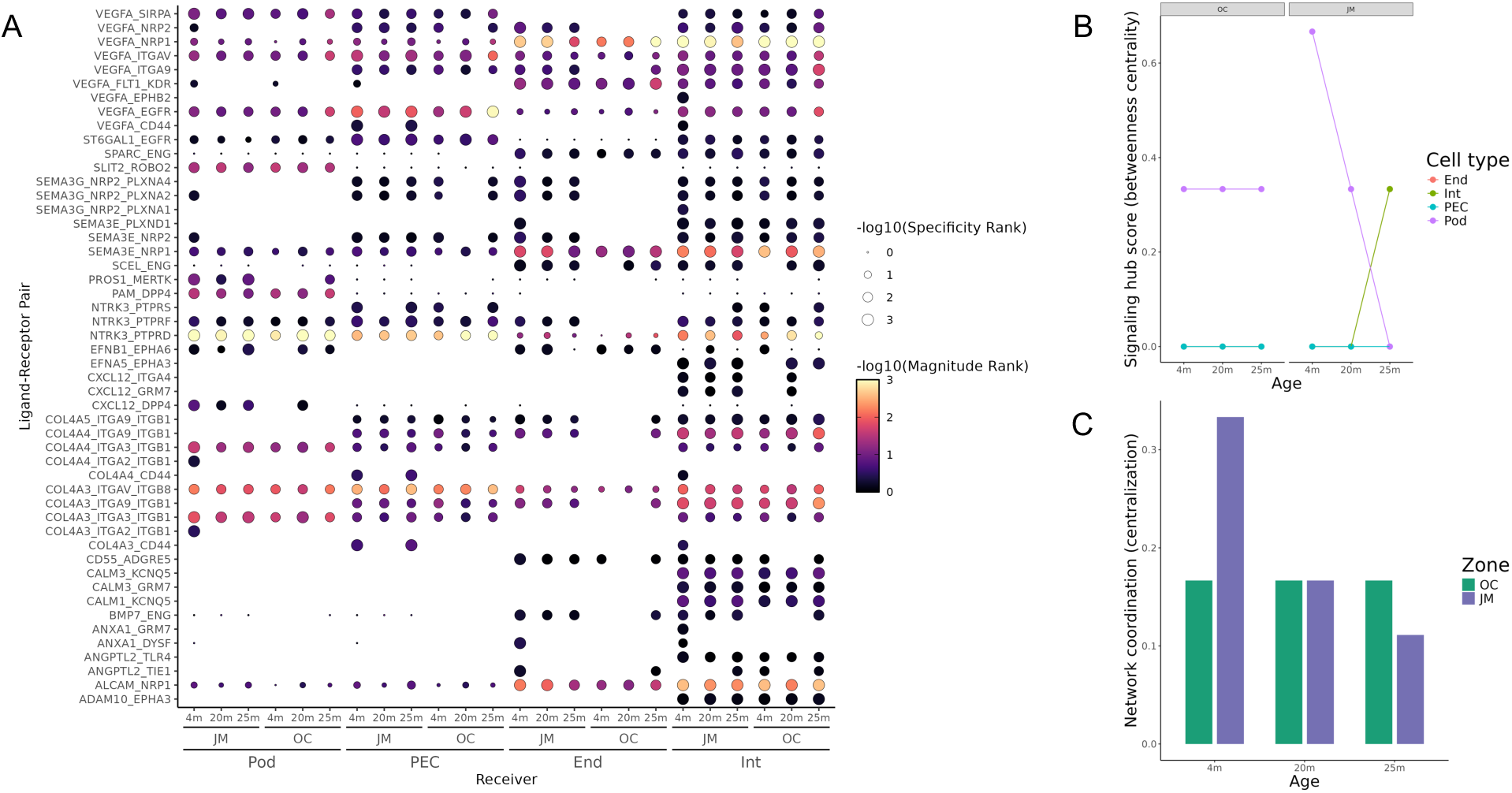
Aging-associated transcriptional remodeling and signaling in podocytes. (**A**) Predicted ligand–receptor signaling from podocytes to all four glomerular cell types, podocytes (Pod), PECs, Mesangial Cells (Int) and glomerular endothelial cells (End), across age and zone. Dot size reflects signaling specificity; color reflects interaction strength, as inferred by LIANA+. (**B,C**) Graph of network topology analysis demonstrating a central role for podocyte-dependent signaling in the young age that persists throughout the ages in the OC but declines in the JM and is replaced by signals from the interstitium (B). This is supported by a bar graph of the network coordination with the OC remaining constant throughout the three age groups, while the JM declining over time (C).

Several ligand–receptor pairs were restricted to specific cellular interactions. For example, podocyte-to-podocyte signaling via Slit2–Robo2 was detected but not observed between podocytes and PECs, mesangial cells, or GEnCs. This pathway has been shown to regulate podocyte adhesion to the glomerular basement membrane.^91,92^ Similarly, Dpp4–PAM interactions were detected specifically within podocyte populations; although not previously characterized in podocytes, inhibition of Dpp4 has been reported to ameliorate renal injury in diabetic models.^93^ In contrast, Adam10–Epha3 interactions were observed selectively between podocytes and mesangial cells, illustrating cell-type-restricted paracrine signaling.

Most signaling relationships were preserved across ages and regions, indicating stable communication architecture. However, a subset of interactions exhibited spatial modulation. Notably, BMP7–Endoglin signaling from podocytes to GEnCs and interstitial cells was detected in the JM at all ages but appeared in the OC only in 25-month kidneys. Because Endoglin mediates TGFβ-driven fibrosis^94^ and BMP7 counteracts TGFβ signaling^91^, this pattern suggests a regionally regulated antifibrotic signaling mechanism.

To determine whether these signaling relationships altered the organization of the glomerular communication network, we quantified network topology using cell-type centrality and global network centralization metrics (**Figure 5B,C**). In young kidneys, JM glomeruli displayed a highly centralized communication structure in which podocytes occupied the dominant hub position. With aging, podocyte hub status progressively declined specifically in JM glomeruli, accompanied by a marked reduction in overall network centralization, whereas OC networks remained comparatively stable. These findings indicate that aging does not merely change signaling activity but reorganizes the communication architecture of the glomerulus, shifting from a coordinated podocyte-centered network to a decentralized multicellular signaling state. Future studies will be needed to demonstrate the functional importance of these.

## DISCUSSION

Kidney aging studies using single-cell and single-nucleus transcriptomic approaches has predominantly focused on tubular compartments—particularly proximal tubules—because of their abundance and susceptibility to injury.^95–102^ In contrast, glomerular aging remains less well defined in unbiased transcriptomic datasets, in part due to the relative scarcity of glomerular cell types, especially podocytes. Here, by profiling more than 890,000 nuclei across age and anatomical regions, we resolved glomerular cell states with sufficient depth to examine both spatial and temporal aspects of aging.

Our findings indicate that kidney aging is not a uniform degenerative process but instead reflects spatially organized and cell-type–specific transcriptional remodeling. Podocytes occupy a structured continuum of states, including a JM-restricted senescent population and intermediate trajectories, while other glomerular cell types remain comparatively stable. Furthermore, ligand–receptor analysis suggests that podocytes act as signaling hubs capable of coordinating multicellular changes within the aging glomerulus. Together, these observations support a model in which glomerular aging arises from reorganization of intercellular regulatory architecture rather than simple accumulation of cellular injury. Podocytes normally function as integrators of multicellular communication, but in aged JM glomeruli this coordinating role is lost, producing a decentralized signaling network. Thus, aging reflects a systems-level failure of tissue regulation rather than independent decline of individual cell types.

In humans, approximately 85% of nephrons are restricted to the OC and 15% are in the JM, differing primarily in glomerular position and loop of Henle length. These structural differences underlie functional specialization, particularly the ability of the kidney to concentrate urine and maintain osmotic gradients. Our analysis demonstrates that this anatomical distinction is also encoded transcriptionally. Using a classifier-based approach, we identified *Kap* and *Napsa* as strong predictors of nephron origin. While *Kap* expression is largely restricted to proximal tubules,^78,79^ Napsa marked JM identity across multiple cell types, including proximal tubules, podocytes, and PECs (**Figure 1G,H**; **Figure 3G**). Although the functional role of Napsa in the kidney remains unclear, its broad and consistent enrichment across JM populations indicates that regional nephron identity is molecularly encoded rather than purely anatomical. This provides a framework for understanding how aging processes can preferentially affect specific nephron populations and supports the concept that spatial context is a determinant of cellular aging trajectories. If regional identity is transcriptionally encoded, aging phenotypes would be expected to emerge preferentially within specific nephron populations rather than uniformly across the kidney.

Regional differences between OC and JM proximal tubules were anticipated, given the anatomical localization of specific segments such as the S3 segment within the JM region (**Figure 2G,H**). In contrast, although morphological and functional differences between OC and JM podocytes have been described, ^88,89,103,104^ the molecular basis for this heterogeneity has remained unclear. Our analysis demonstrates that while the canonical differentiated podocyte population is similarly represented across regions, distinct podocyte subpopulations are spatially biased. Notably, the senescent podocyte state is strongly enriched in the JM region (**Figure 3K**). This suggests that regional microenvironmental conditions, rather than chronological age alone, influence the trajectory of podocyte aging. This spatial bias indicates that vulnerability to aging is dictated by local context rather than time alone. The presence of a JM-restricted senescent population may help explain functional differences between nephron populations, as senescent cells adopt a senescence-associated secretory phenotype (SASP) capable of promoting local inflammation and tissue remodeling.^105^ Importantly, our network analysis indicates that the consequence of this transition is not simply increased inflammatory signaling but loss of centralized control of communication, suggesting that senescence disrupts regulatory coordination within the glomerulus. These findings further raise the possibility that targeting senescence-associated signaling, including emerging senomorphic strategies,^106,107^ could selectively modify regional glomerular aging without broadly affecting all nephron segments.

With respect to aging-associated cellular composition, senescence-associated transcriptional states were detected across multiple renal lineages, but their relationship to chronological age differed by cell type. In proximal tubules, senescence-associated clusters increased with age, consistent with progressive accumulation of injured epithelial cells (**Figure 2I,J**). In contrast, senescent podocyte populations did not increase in abundance in older animals and were present at similar frequencies across all age groups (**Figure 3L**). Rather than contradicting age-dependent decline, this pattern is consistent with the unique biology of podocytes. Senescent podocytes are prone to detachment from the glomerular basement membrane or cell death, limiting their persistence within the tissue and consequently their capture in single-nucleus datasets. Thus, a stable proportion of senescent podocytes likely reflects ongoing turnover rather than absence of aging. This interpretation aligns with established observations that podocyte number progressively decreases with age.^9^ A healthy young human glomerulus contains approximately 500–600 podocytes,^86,108^ declining by roughly 0.9% per year during aging.^40^ Similar rates have been described in rodents, ^25,28,31,109–112^ with region-dependent loss across OC and JM glomeruli.^25^

Thus, aging manifests differently across renal lineages, through accumulation of altered cells in epithelial compartments and progressive depletion in podocytes. A consequence of podocyte detachment from the glomerulus is their appearance in the urine, providing a potential window into podocyte aging in vivo. Viable podocytes can be detected in urine^37^ and individuals over 60 years of age exhibit approximately a 3.3-fold increase in urinary podocytes compared with younger individuals.^40^ Urinary podocyte detection is therefore considered a sensitive indicator of ongoing glomerular injury, and podocyte-specific mRNAs are detectable in the urine of patients with glomerular disease such as FSGS.^113–115^ More recently, single-cell RNA sequencing has been used to characterize the transcriptome of disease-associated urinary podocytes.^116,117^ Our observation that senescent podocytes do not accumulate in tissue but instead likely undergo continual loss suggests that urinary podocytes may capture aging-associated podocyte states that are underrepresented in kidney tissue datasets. If aging podocytes are continuously lost, urinary profiling may preferentially sample late-stage aging states that tissue sequencing cannot easily capture.

During aging, podocyte loss is accompanied by compensatory hypertrophy, in which remaining podocytes enlarge to maintain filtration surface area. Although initially adaptive, hypertrophy becomes maladaptive and is associated with reduced expression of key podocyte genes (*WT1*, *Pod1*, *Nphs1*, *Podxl*), foot process widening, and declining filtration efficiency.^118^ Consistent with this, several podocyte states identified here exhibited reduced expression of canonical differentiation markers. However, we did not identify a transcriptionally distinct hypertrophic population, likely because podocyte hypertrophy is primarily regulated by mTORC1 signaling,^119,120^ which alters translation, autophagy, and apoptosis rather than generating strong transcriptional programs.^121,122^ Together, these observations indicate that structural adaptation, cellular loss, and transcriptional remodeling represent complementary dimensions of podocyte aging. Beyond intrinsic podocyte changes, our data suggest that aging of the glomerulus is coordinated through intercellular communication. Ligand-receptor analysis revealed structured autocrine and paracrine signaling networks centered on podocytes, including interactions selectively directed toward mesangial and endothelial populations. These interactions were largely preserved across age but displayed regional modulation, supporting a model in which podocytes act as signaling hubs that regulate the surrounding glomerular microenvironment. Such signaling provides a plausible mechanism linking localized podocyte state transitions to multicellular remodeling of the aging glomerulus.

In summary, our data support a model in which kidney aging is a systems-level process characterized by spatially restricted collapse of coordinated intercellular regulation. Podocytes normally organize glomerular communication networks, but during aging and particularly in the juxtamedullary nephrons, this hub function is lost, producing decentralized signaling and multicellular remodeling. Thus, aging does not simply alter signaling pathways; it reorganizes the communication topology of the glomerulus. This framework provides a mechanistic explanation for regional susceptibility of nephrons and suggests that therapeutic strategies restoring intercellular coordination may be more effective than those targeting individual pathways. Future studies will be needed to identify, how these changes functionally impact the kidney in general and the podocytes in particular.

## SUPPLEMENTARY FIGURES

**Supplemental Figure 1.**
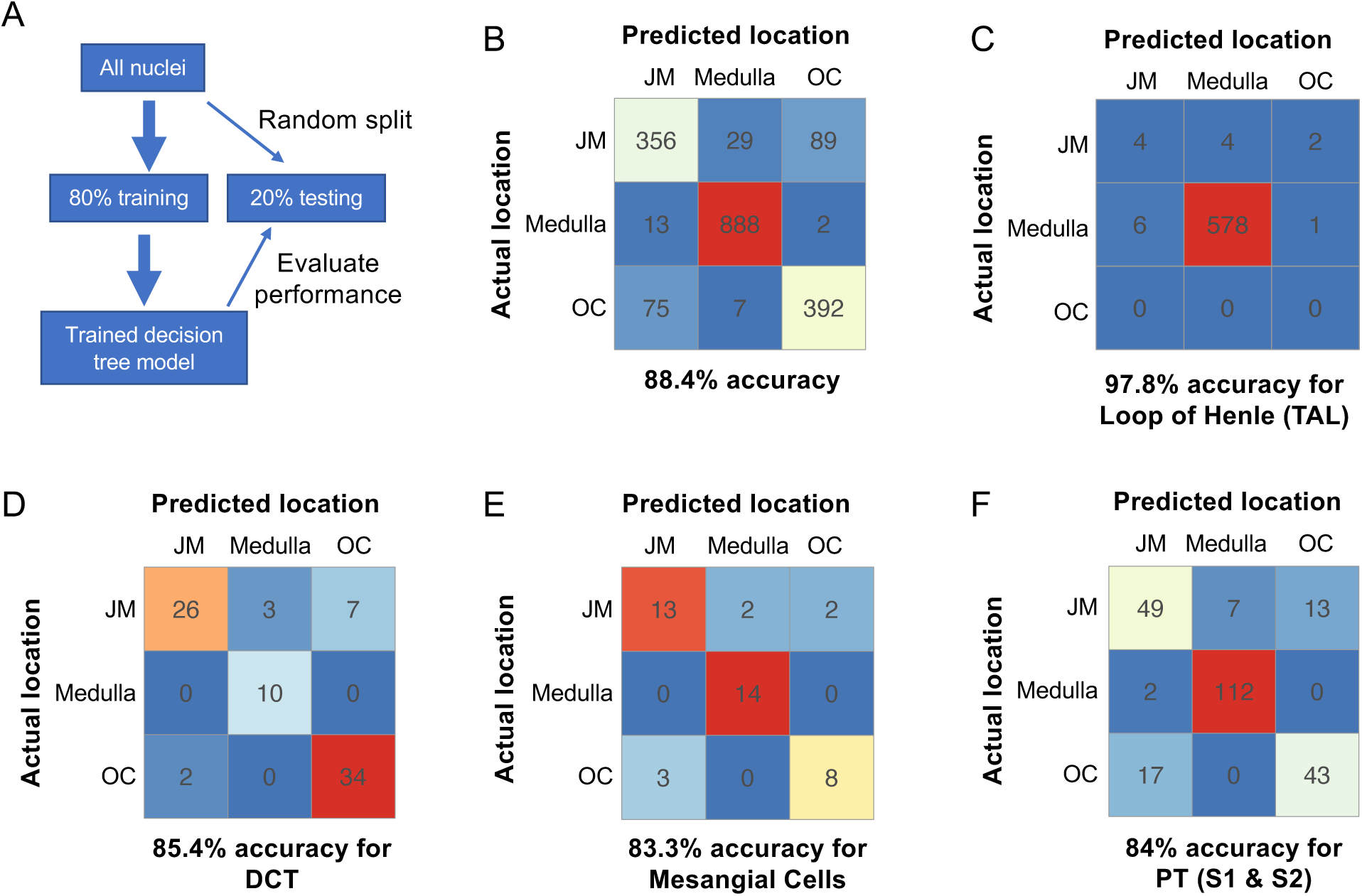
Algorithm to determine spatial expression. (**A**) Schematic of the workflow to accurate assign spatial location from transcriptomic profiles alone training a machine learning model. (**B-F**) Comparison of the accuracy of the model with the actual localization covering all three regions, JM, medulla and OC, with an overall accuracy of 88.4% for spatial location prediction (B). When analyzed by cell type, the prediction accuracy is very high (97.8%) for the TAL, which is complete refined to the medulla (C) and slightly below average for the distal convoluted tubule cells (DCT), mesangial cells and the proximal tubule cells of the S1 and S2 segment (D-F).

**Supplemental Figure 2.**
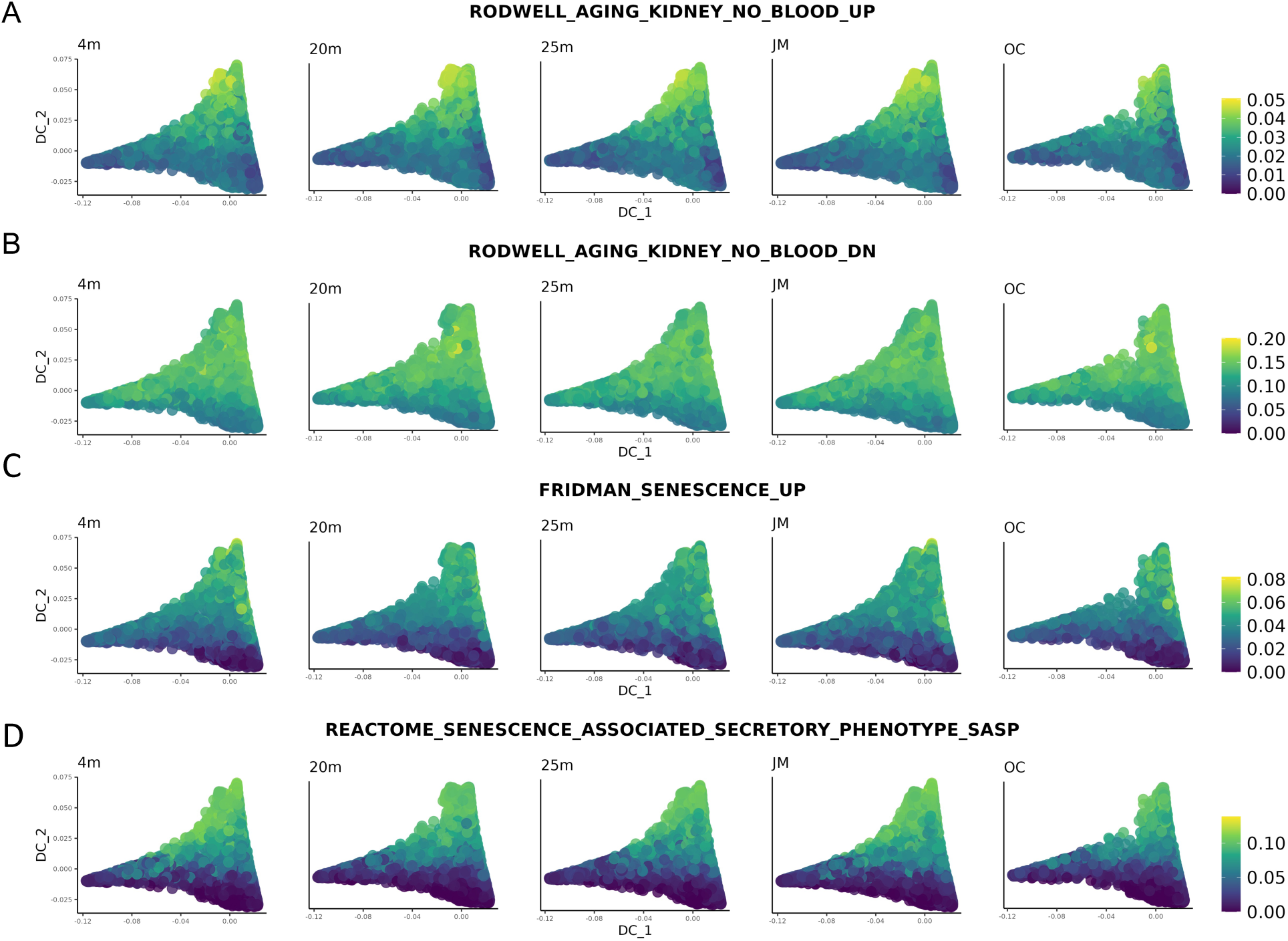
Enrichment of aging and senescence programs in proximal tubule clusters. **(A-D)** Diffusion maps for age and anatomical zone and overlayed with the enrichment scores for the aging and senescence gene signatures Rodwell_Kidney_Aging_Up (A), Rodwell_Kidney_Aging_Down (B), Fridman_Senescence_Up (C), and Reactome_Senescence_Associated_Secretory_Pathway (D) demonstrating enrichment in clusters 1, 7, and 8.

**Supplemental Figure 3.**
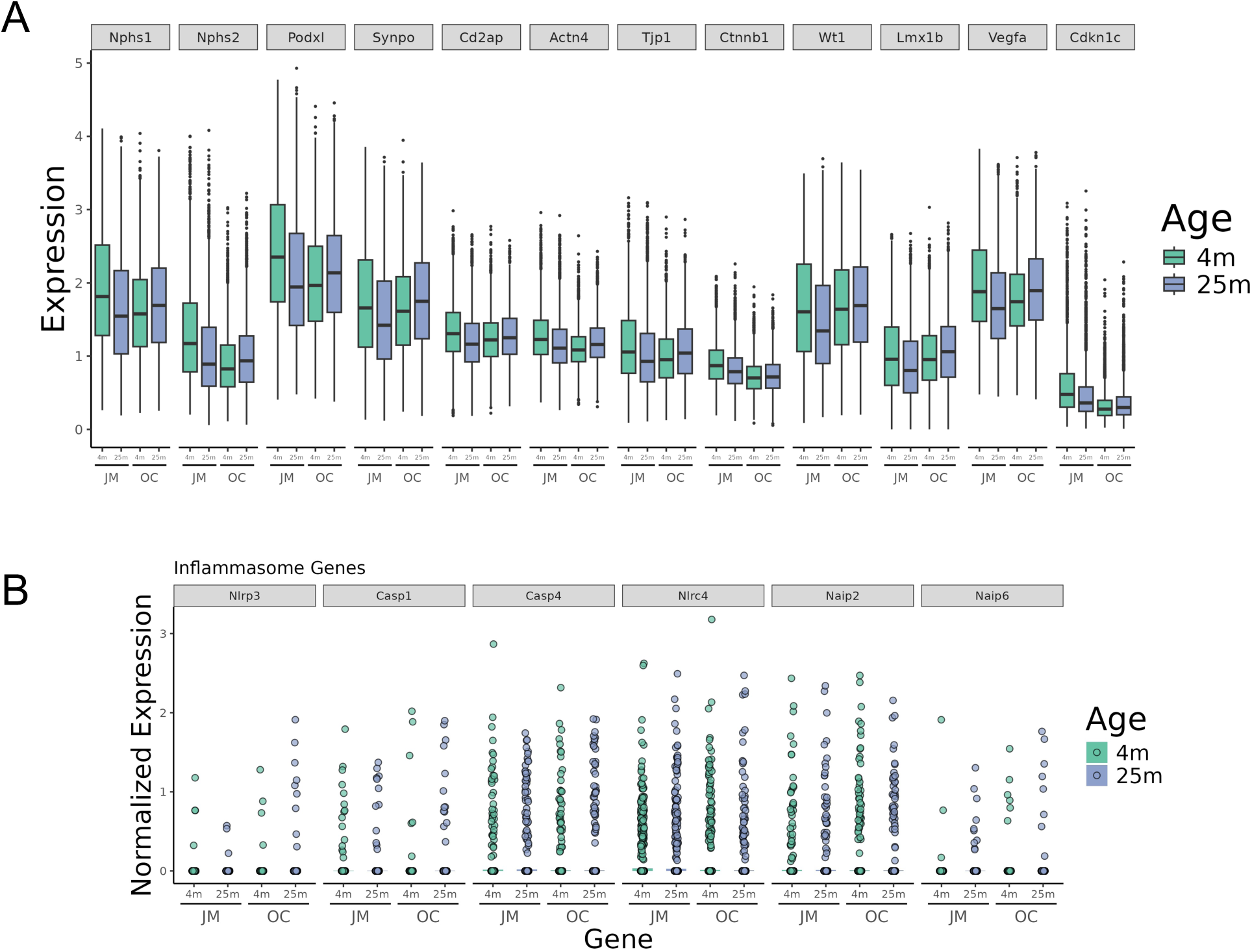
Characterization of canonical and inflammasome gene expression in podocytes. (**A,B**) Expression of canonical (A) and inflammasome genes (B) in podocytes by age (4 months *vs.* 25 months) and zone (JM *vs.* OC). Bars show mean log-normalized expression and overlaid points represent individual nuclei. Note that JM podocytes show reduce expression of canonical genes and elevated expression of inflammatory genes with age.

**Supplemental Figure 4.**
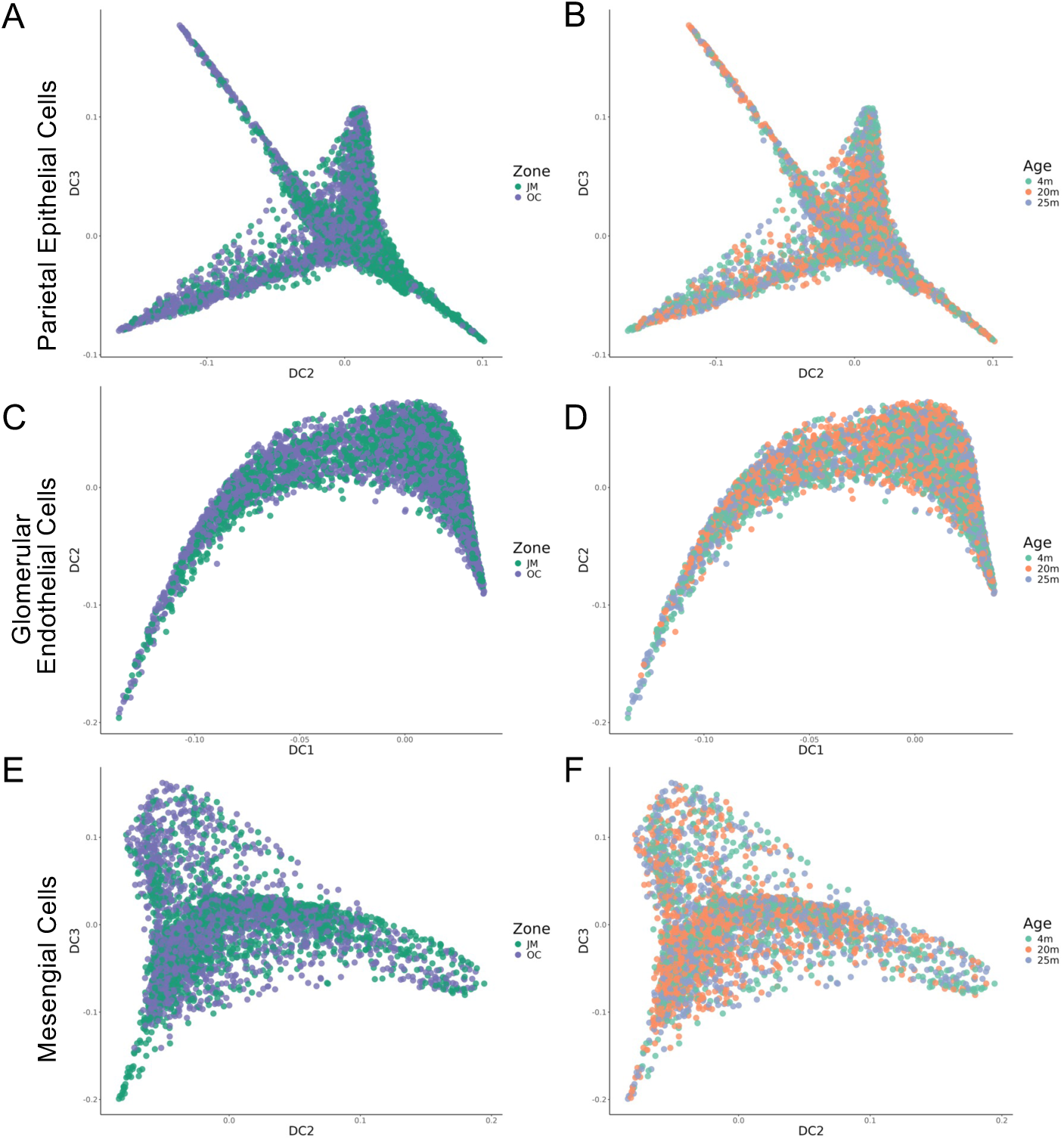
Characterization of non-podocyte glomerular cell types. **(A, D, G)** Diffusion maps for parietal epithelial cells (A,B), glomerular endothelial cells (C,D) and mesangial cells (E,F) overlayed by either anatomical zone (A,C,E) or age (B,D,F) show only little changes in respect to their distribution across the multiple clusters.

