## Supplementary Methods for "SPATIALLY PATTERNED PODOCYTE STATE TRANSITIONS COORDINATE AGING OF THE GLOMERULUS"

**Software Versions and Computational Environment**

All analyses were performed on a Linux-based high-performance computing (HPC) environment.

R analyses were conducted using R version 4.X.X[1] with Bioconductor[2] packages:

DropletUtils (v1.28.0)[3]
scater (v1.36.0)[4]
scran (v1.18.0)[5]
scuttle (v1.18.0)[4]
zellkonverter (v1.18.0)[6]
AUCell (v1.30.1)[7]
slingshot (v2.16.0)[8]
BiocParallel (v1.42.01)[9]

Python analyses were conducted in Python 3.10 using:

scanpy (v1.11.2)[10]
scvi-tools (v1.2.2)[11]
scikit-learn (v1.6.1)[12]

Custom modules:

utils/preprocess.py
utils/model.py
utils/graph_ops.py
eigen-marker

All workflows were executed via scripted pipelines with fixed random seeds.

**Configuration Parameters**

Centralized parameters were defined in config.yaml:

metadata_file: data/metadata/sample_metadata.csv
merged_file: data/processed/merged.h5ad
filtered_file: data/processed/filtered.h5ad
processed_file: data/processed/processed.h5ad
scvi_model_dir: data/processed/model/
batch_key: sample
n_top_genes: 2000
n_latent: 20

Parameters were imported across scripts using the yaml library to ensure consistency.

### Data Filtering and Normalization

Mitochondrial genes were identified using the regex:

^mt-

Cells were removed if they were mitochondrial outliers using:

```
isOutlier(min.diff = 0.5, type = "higher", log = TRUE)
```

Size factors were computed using:

```
quickCluster()
computeSumFactors()
```

Log-normalized counts were generated with:

```
logNormCounts()
```

Highly variable genes were identified using:

```
modelGeneVarByPoisson()
```

The top 10% by biological variance were retained.

#### Conversion to AnnData and Dataset Integration

Each SingleCellExperiment object was converted using:

```
writeH5AD()
```

Datasets were merged using:

```
load_and_merge_h5ad()
```

Genes were filtered using:

```
filter_genes_and_hvg()
```

This retained the top 2,000 variable genes via pp.highly_variable_genes().

### scVI Model Training

The model was trained using:

```
SCVI(adata, batch_key="sample", n_latent=20)
```

Training settings:

Adam optimizer
KL annealing
early stopping enabled

Latent representation extracted using:

get_latent_representation()

Training performed on GPU (NVIDIA Tesla V100).

#### Clustering and Embedding

Neighbor graph:

sc.pp.neighbors(use_rep="X_scVI", n_neighbors=8)

UMAP embedding:

min_dist = 0.0
spread = 2.0
n_components = 3

Louvain clustering:

resolution = 0.1 (global atlas)

Silhouette scores were computed using silhouette_samples().

### Cell Type Annotation

Clusters were mapped to cell types using a manually curated mapping file:

cluster_to_celltype.csv

Canonical markers and expert knowledge were used.

#### Marker Gene Discovery (EIGEN)

Cluster-level discovery:

n_cells = 131072

Cell-type-level discovery:

n_cells = 32768
exclude = ["Rare", "Undetermined"]

Parameters:

expression_threshold = 1.0
min_cells = 3
direction = "up"
block = age + zone

### Gene Set Enrichment Analysis

Gene sets were retrieved from MSigDB{Liberzon, 2015 #995}:

Collections:

H
C2 (KEGG, Reactome, BioCarta, WikiPathways, CGP)

C3 (TFT:GTRD)
C5 (GO:BP)

Filtered to ≥15 genes.

AUCell scores were computed on:

logcounts matrix
diffusion-imputed matrix

Imputation used a modified MAGIC implementation{Van Dijk, 2018 #899} based on a diffusion kernel.

**Trajectory Inference (Podocytes)**

Diffusion map:

k = 8
k_ = 64
Slingshot lineage:
start cluster = 0
terminal clusters = 1,2,3,5
dimensions = first 3 diffusion components

**Cell-Type-Specific Parameters**

| Cell Type | k_ Resolution | Imputation | Trajectory | Downsample |
|---|---|---|---|---|
| Podocyte | 64 0.25 | Yes | Yes | No |
| PEC | 16 0.125 | No | No | No |
| GEnC Stage 1 | 64 0.25 | No | No | No |
| GEnC Stage 2 | 16 0.125 | Yes | No | No |
| Mesangial Stage 1 | 64 0.25 | No | No | No |
| Mesangial Stage 2 | 16 0.125 | No | No | No |
| Proximal Tubule | 32 0.25 | Yes | Yes | Yes |

**Subsampling of Large Populations**

Proximal tubule nuclei (~500,000 cells) were randomly downsampled:

n = 32,768

seed = 42

without replacement

### Reproducibility and Code Availability

All random seeds were fixed.
All processed datasets (.rds, .h5ad) and intermediate outputs are available from the corresponding author.
Code will be publicly released upon publication.
